# Wafer-scale integration of α-quartz thin films towards super high frequency piezoelectric bioNEMS for arbovirus detection

**DOI:** 10.1101/2024.12.26.630366

**Authors:** R. Rathar, S. Ding, D. Sanchez-Fuentes, C. Jolly, C. André-Arpin, R. Desgarceaux, R. Garcia-Bermejo, J. Gazquez, S. Plana-Ruiz, L. Picas, A. Carretero-Genevrier

## Abstract

Micro and nanoelectromechanical systems (MEMS/NEMS), especially piezoelectric resonators, offer a promising strategy for the manufacturing of point-of-care devices providing rapid, sensitive, and field-deployable tests with minimal user training for the diagnostic of viral infections. High-frequency (HF) MEMS/NEMS have the potential for ultrasensible mass-loading devices. Yet, their use for biomedical applications requires challenging manufacturing qualities. Here, we develop a large-scale chemical integration of epitaxial α-quartz (100) thin films on silicon wafers up to 4-inches. This methodology allows the microfabrication of wafer-scale piezoelectric α-quartz/silicon bioMEMS using a recognition layer capable of selectively detecting emerging arboviruses over other viral loads. Using contact-free vibrometry, we show a mass sensitivity of the bioMEMS device of 22.4 pg/Hz in liquid conditions and a Chikungunya virus limit of detection of 9 ng/ml. To reach piezoelectric transduction for compact quartz sensor devices, we develop NEMS resonators at super HF, i.e., 17.8 GHz with a quality factor of 280 which represents a QxF product of 4.98·10^12^. These α-quartz NEMS can reach thicknesses between 100 and 800 nm and lateral dimensions up to 9 mm^2^. Our work opens the door for cost-efficient single-chip epitaxial piezoelectric α-quartz/Si ultrasensitive NEMS sensors manufactured exclusively by soft-chemistry for biomedical applications and many other fields.

## 1. Introduction

The α-quartz, with a vast resonance quality factor (QxF > 10^13^)[1] and exceptional temperature stability, is the best piezoelectric sensing material. As electric oscillators, quartz single crystals or plates can provide a stable clock frequency in virtually any electronic application. Furthermore, their resonant frequency’s sensitivity to mass and thickness also makes them ideal for use in microgravimetric devices for sensing applications[2]. Therefore, in 2030, an estimated 15 billion quartz crystal resonators and oscillators will be manufactured and placed in automobiles, digital cameras, industrial equipment, etc. Today, most current research on α-quartz focuses on the performance enhancement of single crystals or bulk quartz devices based on top-down strategies[3]. However, fully exploiting the potential of α-quartz and achieving enhanced device performances in future applications requires controlling and scaling up its epitaxial integration on silicon under a thin film form down to the nanoscale. Moreover, the successful wafer-scale implementation of epitaxial quartz thin films on Si is necessary for industrial processing using microelectronic technologies[4]. As a result, the silicon substrate must be chemically and structurally compatible with the α-quartz structure to prevent undesirable interfacial defect formation and be economically feasible for large-scale production. Despite the importance of quartz in leading-edge applications, its industrial integration into silicon has yet to be achieved.

α-quartz crystals applied to microelectronics are exclusively synthesized by hydrothermal methods, making it impossible to reduce the wafer thickness below 30 µm and directly integrate epitaxial quartz on silicon[5,6]. Because the mass sensitivity of thickness shear mode quartz resonator-based devices is inversely proportional to the square of the thickness of the resonator, the wafer thickness is among the critical barriers to overcome towards higher frequency resonators and, thus, transducers with improved sensitivity[7]. Furthermore, the inability to integrate and scale up α-quartz onto silicon wafers represents a significant drawback in achieving improved developments in the α-quartz industry. Consequently, current α-quartz devices can only be assembled through direct bulk micromachining or hybrid integration methods. Therefore, the performance of these sensing devices appears reduced due to the difficulty of preparing devices directly on silicon single crystal substrates or the use of bonding techniques. This criterion is crucial in the context of biological sensing applications using micro and nanoelectromechanical systems (bioMEMS/NEMS), which are subjected to very low-quality factors (1–10) due to the viscous damping. Furthermore, bioMEMS/NEMS must be robust and portable with minimal sample preparation and a multiplex approach allowing access to precise molecular signatures of biomarkers[8]. Despite these technological needs, today, ultrasensitive high-frequency on-chip quartz piezo-bioNEMS devices are not yet developed.

The development of bioMEMS/NEMS as point-of-care (PoC) testing devices provides a strategy for rapid, sensitive, and compact diagnostic solutions with a minimal user training, as opposed to sophisticated clinically-based systems (e.g., RT-PCR or ELISA) that are virtually impossible to deploy in the field and regions with limited resources[9,10]. In this context, the global COVID-19 pandemic and the current drastic rise in the emergence of arthropod-borne viruses (arbovirus) such as Dengue, Chikungunya, West Nile, or Zika virus in large geographic areas highlight the importance and need of conceiving early detection systems to help contain virus spread and infectivity. The integration of piezoelectric materials in MEMS/NEMS for PoC diagnostics offers significant advantages in terms of sensitivity, miniaturization, energy efficiency, and versatility[11–13]. Consequently, developing ultrasensitive piezoelectric bioNEMS should open the door to more efficient, accurate, and sensitive PoC virus detection tests.

In this direction, we previously succeeded in engineering piezoelectric α-quartz thin films on silicon and SOI substrates with controlled microstructure using chemical solution deposition (CSD)[14,15]. This chemical method, which relies on the thermal devitrification of dip-coated mesoporous silica films, assisted by alkaline earth cations in amphiphilic molecular templates has been used, for instance, to develop a biocompatible nanostructured piezoelectric MEMS with a mass resolution down to 100 ng/Hz in air[16,17].

A prerequisite to enable the technological application of piezoelectric resonators based on quartz thin films, for instance, in bioNEMS, is gaining a greater control over the large-scale integration of epitaxial quartz and its microfabrication at the silicon wafer scale. Yet, a method to prepare large scale quartz thin film on silicon has not been reported so far. Furthermore, the synthesis of epitaxial α-quartz thin films by dip-coating has the inconvenience of having an inhomogeneous crystallization at the scale of the silicon wafer, due to a less adapted solution deposition process to the microelectronics standard. This process involves the dip-coating of a silica precursor solution doped with Sr^2+^ devitrifying agent and the appearance of a meniscus, which encompasses thickness variation and layer uniformity issues. Moreover, for most technological applications, it is required to control the crystallization for single chip devices processes at the wafer scale, but the maximum size of epitaxial quartz films prepared by dip coating technique cannot exceed a few cm[18].

In this work, we address these shortcomings by controlling the nucleation, crystallization, microstructure and thicknesses of epitaxial α-quartz films on silicon up to 4-inches[19]. We investigated (i) the quartz precursor solution stability by optimising our precedent chemical formulation, (ii) the deposition conditions using spin-coating technique, which is more adapted to commercial silicon wafer dimensions, and (iii) the crystallisation of epitaxial quartz thin films on 2-, 3- and 4-inches silicon wafers while tailoring the film microstructure and thicknesses. We exploited this wafer scale integration of epitaxial quartz thin films on silicon to fabricate wafer-scale piezoelectric bioMEMS and NEMS high quality factor resonators at super high-frequency, from 5 KHz to 17.8 GHz. We designed a recognition layer based on the ultrastable SpyCatcher/SpyAvidin system for the immobilization of commercial biotinylated antibodies, hence rendering quartz MEMS functional for the selective detection of arbovirus. Using an optical interferometric set-up with an integrated microfluidic system and the Chikungunya virus (CHIKV) as a model of viral sample, we show that quartz bioMEMS exhibit a mass sensitivity of 22.4 pg/Hz in a liquid medium with a limit of detection (LoD) of the CHIKV virus of 9 ng/ml. Finally, we show the development of a piezoelectric NEMS based on a 120 nm-thick quartz membrane resonator with a piezoelectric resonance frequency of 17.8 GHz and quality factor about 280 at air, which correspond to a Qxƒ product of 4.7 10^12^ at 300K, not far from quartz bulk Qxƒ product limit (10^13^)[1]. Collectively, this work shows that by performing measurements at higher frequencies and exploiting the quartz piezoelectric transduction, it should be possible to reach higher sensitivities and implement these performances on a portable PoC device for a wide number of biomedical applications.

## 2. Wafer-scale integration of epitaxial (100) α-quartz thin films on silicon

In this work, we aimed to develop a method to scale up the integration of α-quartz thin films in order to manufacture piezoelectric bio-MEMS/NEMS for the selective detection of arbovirus, as schematized in **Figure S1**. To this end, we first developed an industrial solution based on the ability to scale up, i.e., to integrate quartz-α in epitaxial thin film form on larger surfaces (up to a 4-inch silicon (100) wafer) without compromising the crystal quality, orientation, homogeneity and piezoelectric properties of the material. To accomplish this challenge, we first modified our previous quartz precursor solution and the deposition methodology[15,18] using a spin-coating technique, which appears more adapted to the microelectronics format.

A detailed analysis of the effect of surfactants on strontium catalyst salt solubility was carried out to achieve higher strontium concentrations, thereby improving quartz crystallization as part of the scaling-up of the integration process. In this regard, we incorporated into the α-quartz precursor sol-gel solution Brij-58, a non-ionic surfactant, to increase the solubility of the strontium chloride hexahydrate (SrCl₂·6H₂O) catalyst while preserving the stability of the precursor solution. We found that Brij-58 promotes a better homogeneous distribution of Sr^+2^ devitrifying cations through an evaporation-induced self-assembly (EISA) mechanism[20]. This feature could be induced by the fact that surfactant Brij-58 can form nanometric micelles capable of absorbing small amounts of water[21]. Therefore, these Brij-58 micelles could form during the EISA process of spin-coated doped-strontium silica layers, trapping small quantities of water and Sr^+2^ ions. As a consequence, the molar ratio of Brij-58 (R_Brij_ = 0.044) proved to stabilize strontium salt concentration up to R_Sr_ = 0.1 into quartz sol-gel precursor solution (see **Table S1**), achieving homogeneous nucleation and optimum crystallization of the epitaxial (100) quartz thin films at the wafer scale as shown in **Figure 1a**.

**Figure 1.**
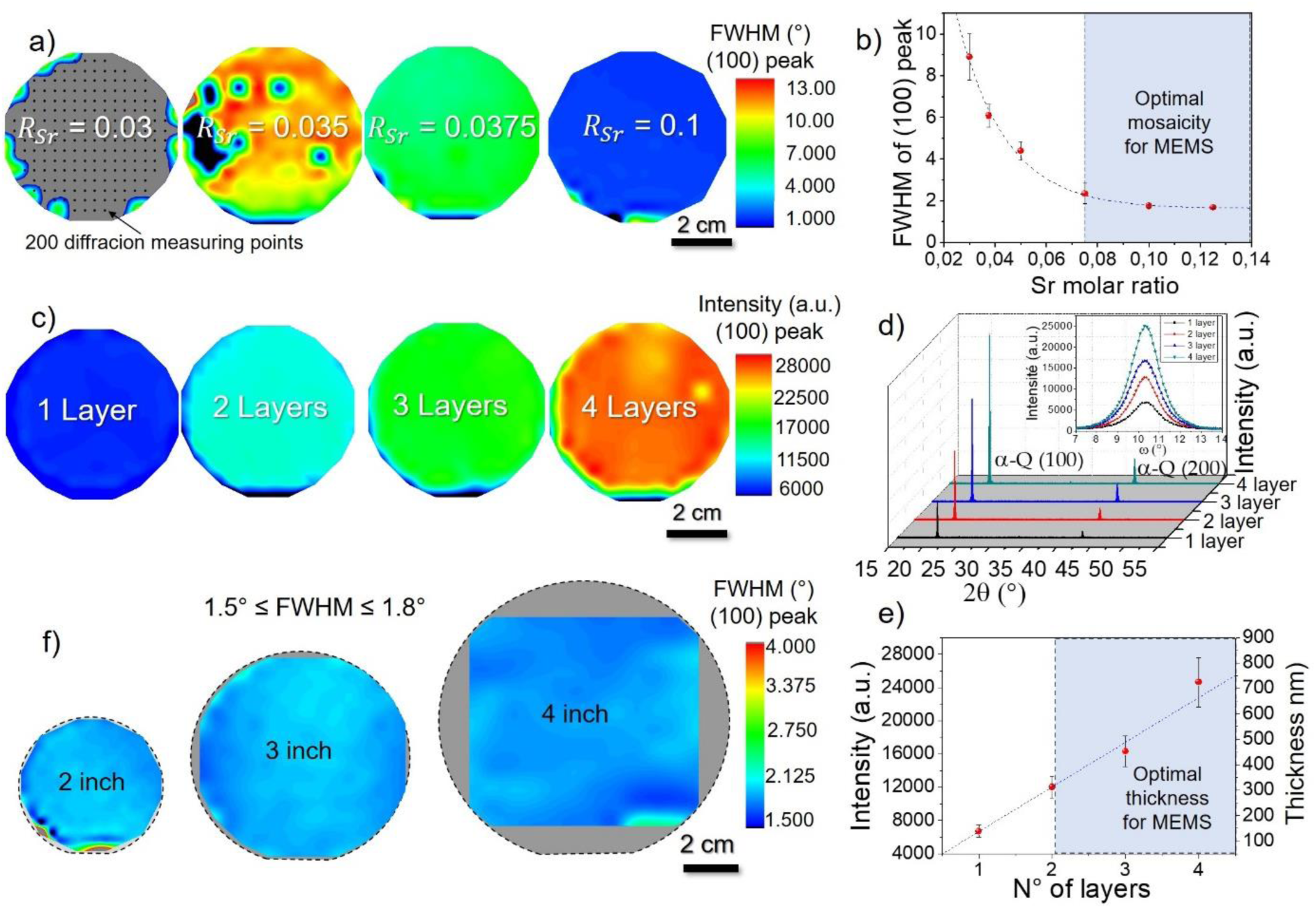
Wafer-scale integration of epitaxial (100) α-quartz thin films on (100) silicon up to 4 inches. **a)** Nucleation and crystallization of epitaxial (100) α-quartz layer on a 2-inch wafer as a function of Sr devitrifying agent molar ratio. Mosaicity heat maps around the (100) α-quartz peak indicate homogeneous nucleation from a Sr ratio of 0.0375. Notice that each textural heat map is composed of 200 diffraction measuring points performed with a collimator of 1 mm^2^ and a 2D XRD detector in order to perfectly cover the whole silicon wafer surface **b)** Plot showing the decrease in FWHM value around the (100) quartz peak as a function of Sr molar ratio. **c)** Intensity heat maps of the (100) quartz peak on 2-inch wafers with different numbers of deposited layers. **d)** XRD θ–2θ scan results for films with different layer numbers. Insets show the FWHM for each sample. **e)** Graphic showing the linear relationship between the (100) quartz peak intensity, number of deposited layers, and total quartz layer thickness. **f)** Mosaicity mappings around the (100) quartz peak, demonstrating the homogeneity of quartz layers on 2-inch, 3-inch, and 4-inch wafers.

For α-quartz thin films prepared with an SrCl₂·6H₂O molar ratios between 0.030 ≤ R_Sr_ ≤ 0.035, the strontium concentration is not sufficient to devitrify the amorphous sol-gel and silicon native SiO_2_ layer and thus impeding quartz nucleation and epitaxial crystallization. However, we found that silica layers doped with an SrCl₂·6H₂O molar ratios greater or equal to R_Sr_ ≥ 0,0375 crystallize uniformly over the entire 2-inch Si wafer substrate (**Figure 1**). More importantly, with a molar ratio R_Sr_ = 0.1, the epitaxial (100) α-quartz layers reduce their degree of mosaicity, reaching uniform 2-inch (100) quartz/ (100) silicon with a top mosaicity degree of 1.13° (**Figure 1b** and **Figure S2.**).

Spin-coating deposition conditions of sol-gel precursor solution also influence the crystallization of quartz layers. We have carefully examined the impact of centrifugation speed on the thickness of the deposited silica layer and the crystallinity of the final epitaxial quartz thin film. As a result, samples manufactured with a centrifugation speed between 1000 < ω < 3500 rpm and a fixed optimized surfactant molar ratio of R_Brij_ = 0.044 and R_Sr_ = 0.1 were prepared (**Figure S3**). We observed that silica films deposited with a centrifugation speed of ω = 2000 rpm for 30 seconds obtained 220 nm thick uniform silica layers with the best mosaicity degree at the silicon wafer-scale after crystallization at 1000 °C under an air atmosphere. Taking advantage of all these results, we were able to fabricate uniform and flat thin films of epitaxial (100) quartz at the wafer scale optimal for MEMS applications with small thickness variations in the 170 nm to 285 nm range (see **Figure 1b** and **Figure S4**).

To achieve a larger and controlled thickness of epitaxial quartz thin films at a wafer scale, we have developed a multi-deposition approach by using spin-coating (**Figure 1c**). This methodology consists of depositing several strontium-doped silica sol-gel layers pre-consolidated at 500 °C. After the quartz crystallization process, this multi-layer deposition method showed a linear increase of the layer thicknesses up to 710 nm while maintaining the crystal quality, mosaicity, and coating uniformity without the appearance of secondary phases or undesirable quartz orientations (**Figure 1d** and **Figure 1e**). Finally, thanks to the optimum conditions for integration of quartz on Si, we carried out the crystallization of strontium-doped silica layers deposited on 2, 3- and 4-inches silicon wafers. Hence, these results demonstrate the scaling of the integration process. Furthermore, our results showed an equivalent crystallization irrespective of the substrate size, indicating homogeneous nucleation across the entire substrate with mosaicities between 1.5° and 1.8° and free of macroscopic defects (**Figure 1f** and **Figure S5)**.

## 3. Epitaxial characterization of (100) α-quartz thin films wafers on (100) silicon

Next, we studied the epitaxy of α-quartz films with three-dimensional electron diffraction (3D ED) in film cross-section, which is a technique that enables the reconstruction of the diffraction space from nanometre-sized regions[22]. We collected two 3D ED datasets with a 120-nm electron beam just above and below the interface during a single tilt scan of the TEM goniometric stage (**Figure 2a**). The acquired diffraction data was processed to determine the unit cell parameters of the silicon and quartz areas and understand its crystallographic relation. The parameters found for quartz (a=b=4.9448(3) Å, c=5.4261(2) Å, α=β=90.0°, γ=120.0°) confirmed the presence of the α polymorph and a dynamical refinement with electron precession allowed the determination and refinement of the corresponding absolute crystal structure (**Table S2.**). The retrieved orientation matrices showed that the ***a*** and ***c*** vectors of quartz lay on the interface plane with a slight misalignment to the respective vectors of Silicon 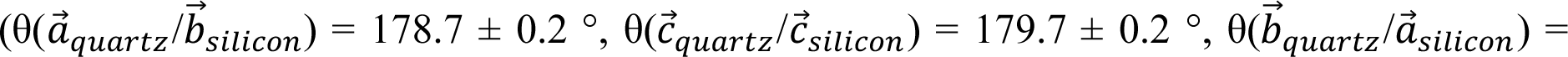 61.3 ± 0.2 °). Interestingly, the ratio between *b⃗_silicon_* and *a⃗_silicon_* is around 10%, which means that the initial match of the unit cells is recovered after 10 cells. Although such high displacement of the α-quartz is significant, it does not seem to produce substantial defects on the epitaxy as confirmed by atomic resolution bright field (BF) STEM imaging (**Figure 2b**), in which a sharp interface is observed. Pole figures of the Si{100} and α-quartz {101} reflections of **Figure 2c** confirmed the previously observed epitaxial growth and the existence of two well-defined quartz crystal domains that have the same epitaxial relationship of [100]Si(100)//[100]*α-quartz(100)*, in agreement with our previous report[15].

**Figure 2.**
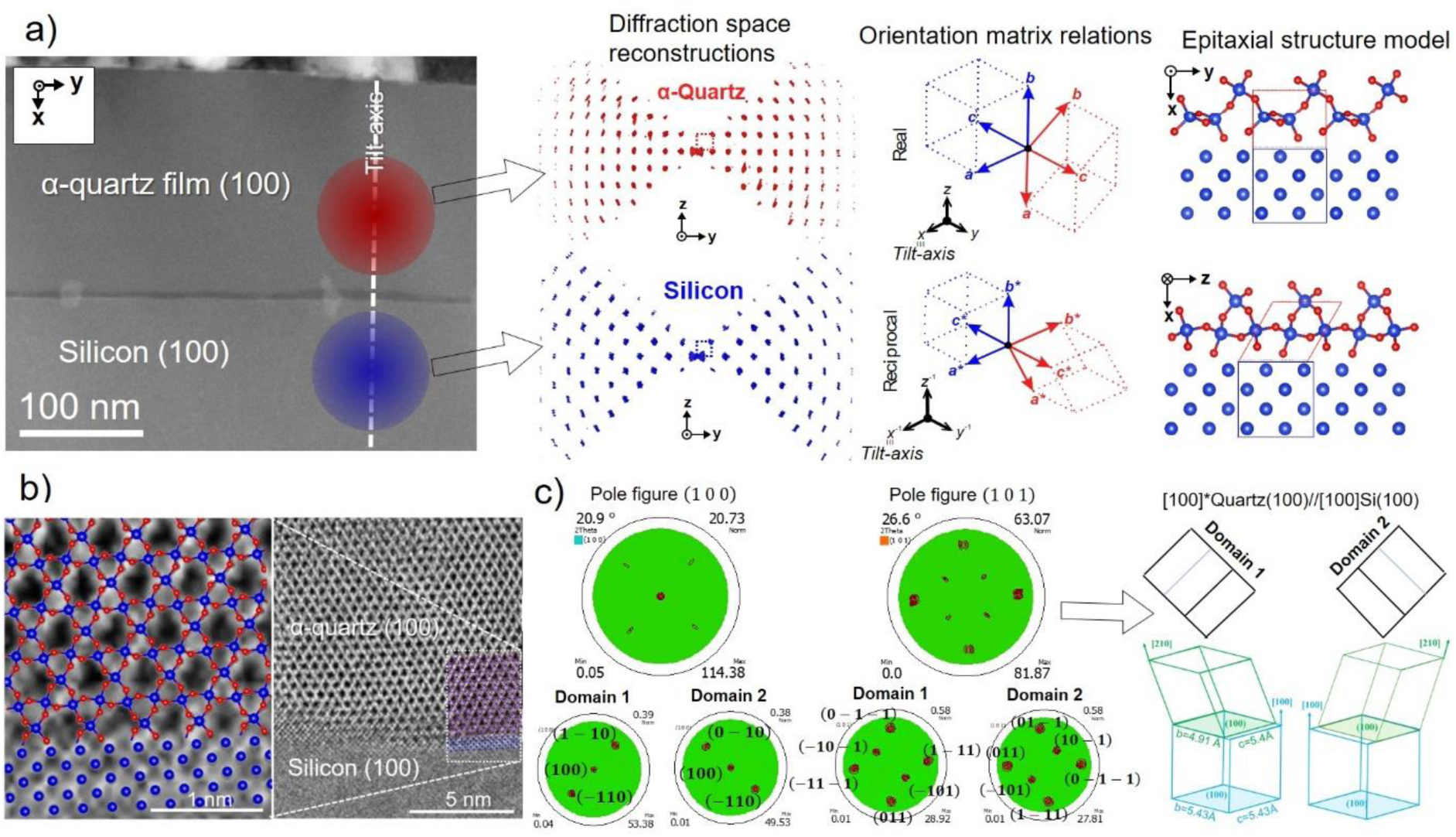
Characterization of epitaxial (100) α-quartz thin films on (100) silicon wafer. **a)** Three-dimensional electron diffraction analysis in quartz film cross-section. **b)** Atomic resolution BF images of the (100)α-quartz/(100)Si interface acquired in STEM mode. **c)** Indexed pole figures of Si{100} and α-quartz {101} reflections that demonstrate the existence of two well-defined quartz crystal domains as schematically represented in 3D on the right side of the figure.

## 4. Bio-conjugation of epitaxial α-quartz wafers on silicon with a virus-recognition layer

Next, we set up a bio-conjugation system to use α-quartz thin films as a piezoelectric bioMEMS for the selective detection of viral loads (**Figure 3**). Indeed, the conjugation of a recognition layer in the α-quartz thin film is essential to give bioMEMS specificity in the detection of a biomarker of interest (e.g., a specific virus)[23]. To this end, we used a covalent strategy based on the silanization of α-quartz films hence, allowing the functionalization of its surface with alkoxysilane molecules. To this first layer we covalently conjugated a layer of the SpyCatcher protein, which forms a spontaneous isopeptide bond with SpyTag hence forming an efficient and irreversible SpyTag/SpyCatcher complex[24,25]. We set up the concentration of SpyTag and SpyCatcher allowing a proper surface coverage of α-quartz surfaces by using a fluorescently tagged version of each part of the complex, namely SpyTag-mKate2 and SpyCatcher-GFP[26], and evaluate their surface association by confocal microscopy (**Figure 3a-e**). As shown in **Figure 3a-b**, we observed a concentration-dependent increase of the fluorescence intensity of the SpyCatcher-GFP signal upon bio-conjugation of the α-quartz surfaces after 1h of reaction time followed by several rounds of rinsing with a phosphate buffered saline (PBS) solution. Hence, indicating a permanent and robust conjugation between the aminosilanized surface and SpyCatcher. Furthermore, we found that, in the absence of SpyCatcher, SpyTag cannot associate with APDMES-treated surfaces passivated with bovine serum albumin (BSA), even upon increasing the SpyTag concentration (**Figure 3c**), pointing out that SpyCatcher is necessary for SpyTag conjugation in our system. Increasing concentration of SpyCatcher led to a parallel increase of the SpyTag-mKate2 intensity, confirming the formation of the isopeptide and the covalent tethering of the SpyTag/SpyCatcher complex on the α-quartz surface (**Figure 3d**). Importantly, fluorescence analysis was evaluated after serval rounds of extensive rinsing with PBS, confirming the stability of SpyTag/SpyCatcher bio-conjugated surfaces. Moreover, the assessment of the fluorescent images showed a homogenous functionalization of α-quartz surfaces at different ratios of the SpyTag/SpyCatcher complex (**Figure 3e**).

**Figure 3.**
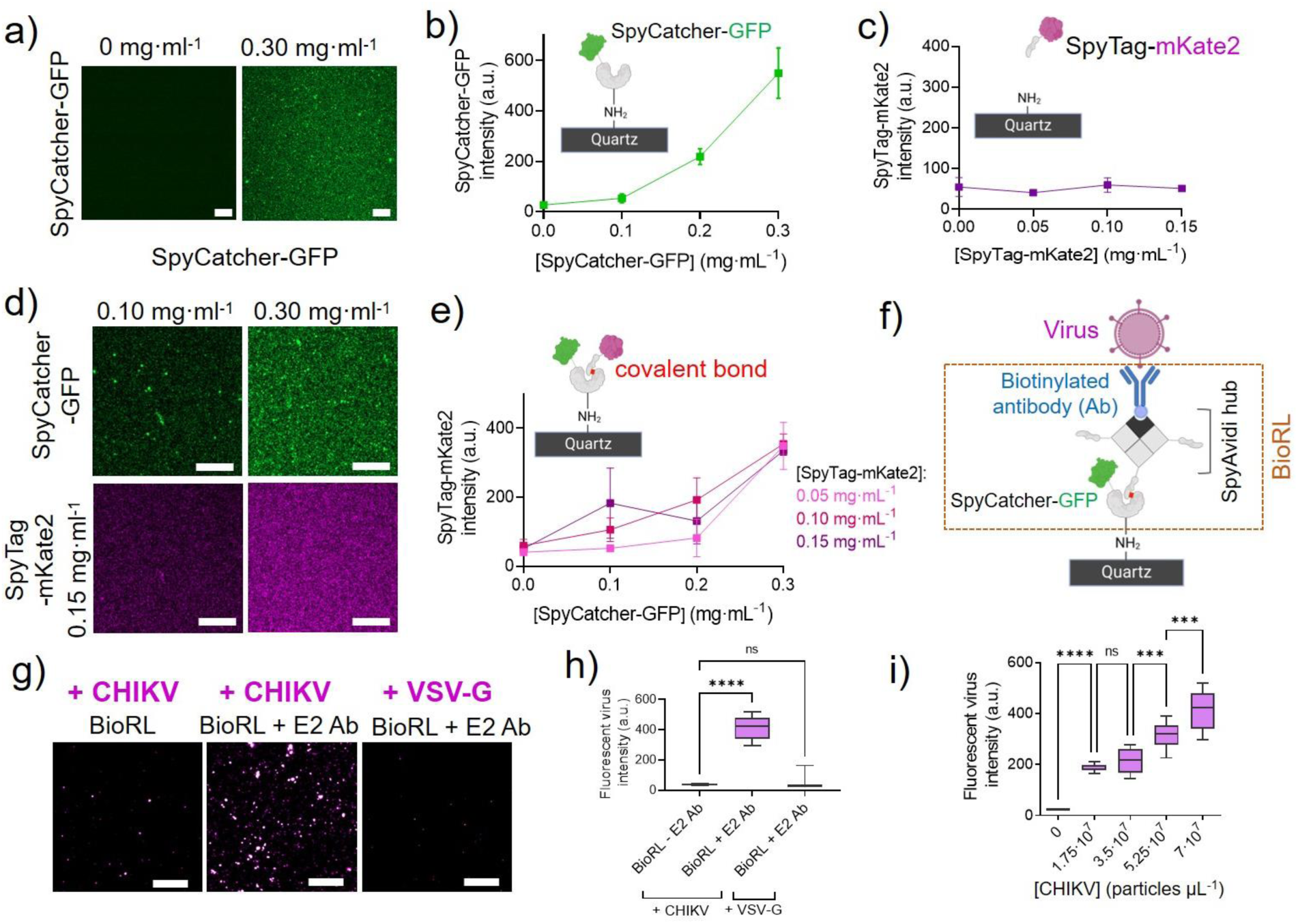
Bio-conjugation of a SpyCatcher/SpyAvidin virus-recognition layer (bioRL) on α-quartz thin films. **a**) Confocal image of the surface conjugation of the SpyCatcher-GFP (green) layer at two concentrations: 0 mg·mL^-1^ (background condition) and 0.30 mg·mL^-1^. Scale bar, 10 μm. **b)** Intensity of the SpyCatcher-GFP layer at different concentrations to SpyCatcher-GFP. Mean ± s.d. **c)** Intensity of SpyTag-mKate2 association α-quartz surfaces at different concentrations in the absence of SpyCatcher. Mean ± s.d. **d)** Confocal images showing the surface conjugation of the SpyCatcher-GFP (green): SpyTag-mKate2 (magenta) complex at two different ratios, i.e., (1:1.5) and (2:1), respectively. Scale bar, 10 μm. **e)** Intensity of SpyTag-mKate2 at different concentrations as a function of the concentration of SpyCatcher-GFP conjugated layer. Mean ± s.d. **f)** Schematic representation of the bio-recognition layer (bioRL) for virus detection (in magenta) using the SpyCatcher/SpyAvidin complex consisting of a tetramer of three dead streptavidin fused to SpyTag (in gray) forming an isopeptide bond with SpyCatcher-GFP and one traptavidin (in black) binding to a biotinylated antibody (in blue). **g)** Confocal images showing fluorescent virus-like particles isotyped with the envelop proteins of CHIKV (+ CHIKV) or VSV-G (+ VSV-G) attached to the bioRL in the absence (bioRL – E2 Ab) or presence (bioRL + E2 Ab) of a biotinylated antibody against CHIKV E2 proteins. Scale bar, 10 μm. **h)** Box plot of the intensity of fluorescent virus particles interacting with the bioRL in the absence (bioRL – E2 Ab) or presence (bioRL + E2 Ab) of a biotinylated antibody against CHIKV E2 proteins. Mean ± s.d. **i)** Box plot of the intensity of fluorescent CHIKV particles attached to the bioRL from samples loaded with different concentrations of viral particles. Mean ± s.d. Statistical differences were assessed by Mann-Whitney test: n.s > 0.1, ***P < 0.001, ****P < 0.0001. Data represents 6 experimental replicates. Schematics were created with BioRender.com.

To render the SpyTag/SpyCatcher-based recognition layer selective to a specific virus, we set up a strategy based on the capacity of commercial antibodies to specifically detect the structural proteins of viral particles, such as the envelope (E) glycoproteins. Indeed, envelope proteins have been extensively used as a target for an accurate diagnostic tool[27], since they are highly specific to a given virus and even, between variants of the same virus[28]. To this end, we used SpyAvidin hubs[29], which consist of SpyTag-traptavidin subunits with maximized biotin binding strength allowing the ultrastable orthogonal assembly of a single biotinylated antibody on each chimeric SpyAvidin tetramer (**Figure 3f**). As a proof-of-concept for the bio-recognition layer (bioRL) system, we set out to detect the Chikungunya virus (CHIKV) using a commercial mouse monoclonal biotinylated antibody recognizing the E2 protein of CHIKV. To this end, we generated Chikungunya virus (CHIKV) pseudo-typed particles incorporating the E1 and E2 envelope proteins from the CHIKV LR2006 OPY-1 variant, hereafter called CHIKV particles, as described in the methods section. Furthermore, to facilitate the validation of the bioRL by fluorescence microscopy, we generated a fluorescent version of CHIKV particles through the fusion of E1 to an mRuby3 tag. As shown in **Figure S6a**, we found a perfect co-localization between the anti-E2 mouse monoclonal antibody and fluorescent CHIKV particles spotted on α-quartz surfaces, confirming the specificity of the antibody to be further used for the bioMEMS bioRL. We then assessed by fluorescence microscopy the ability of the bioRL to capture CHIKV particles (**Figure S6b-c**). We found that CHIKV particles are captured by the bioRL only when the anti-E2 antibody is conjugated at a ≥ 1:0.5 ratio of the SpyCatcher/SpyAvidin complex.

Next, we assessed the selectivity of the bioRL over different viral samples. To this end, we generated fluorescent pseudo-typed viral particles with the envelope glycoproteins of the vesicular stomatitis virus G (VSV-G particles). We analyzed by airyscan/confocal microscopy the binding of CHIKV and VSV-G particles to the anti-E2 antibody conjugated bioRL and found a significant capture of CHIKV particles over the other viral sample (**Figure 3g-h**). Furthermore, we observed a linear increase of the fluorescent virus intensity captured by the bioRL with increasing concentrations of CHIKV particles. Altogether, confirming the selectivity of the engineered bioRL to detect CHIKV loads over other viral samples.

## 5. Wafer-scale microfabrication and electromechanical characterization of piezoelectric α-quartz/Silicon MEMS

The successful wafer scale integration of high-quality epitaxial α-quartz thin films on silicon has led us to the engineering of quartz-based microelectromechanical (MEMS) device sensor at the wafer scale with control over resonant frequency, dimensions, thickness, and morphology as shown in **Figure 4a**. We decided to fabricate membrane resonators because they are better adapter for biosensor applications, especially in the environmental and biomedical/pharmaceutical fields[30]. To this end, we developed a cost-effective microfabrication methodology combining laser and chemical etching on the scale of a single 2-inch silicon wafer[31]. This microfabrication approach combines a direct laser engraving process with a selective anisotropic wet etching with tetramethylammonium hydroxide (TMAH) from the backside of the silicon substrate (see **Figure S7**). The laser etching process allowed the desired design of the aperture and, thus, the final shape of the membrane resonator. Moreover, the laser etching step process drastically increased the speed of silicon perforation in the initial cavity, resulting in a much faster process than by chemical etching. (see **Figure S6**). To protect the backside silicon wafer from the chemical etching, we developed an innovative layered system of protective coatings made 400 nm thick SiN /400 nm thick SiO_2_/ 400 nm thick SiN which made it possible to resist the chemical attack of TMAH for more than 20 hours at 84 °C. (see **Figure S8**).

**Figure 4.**
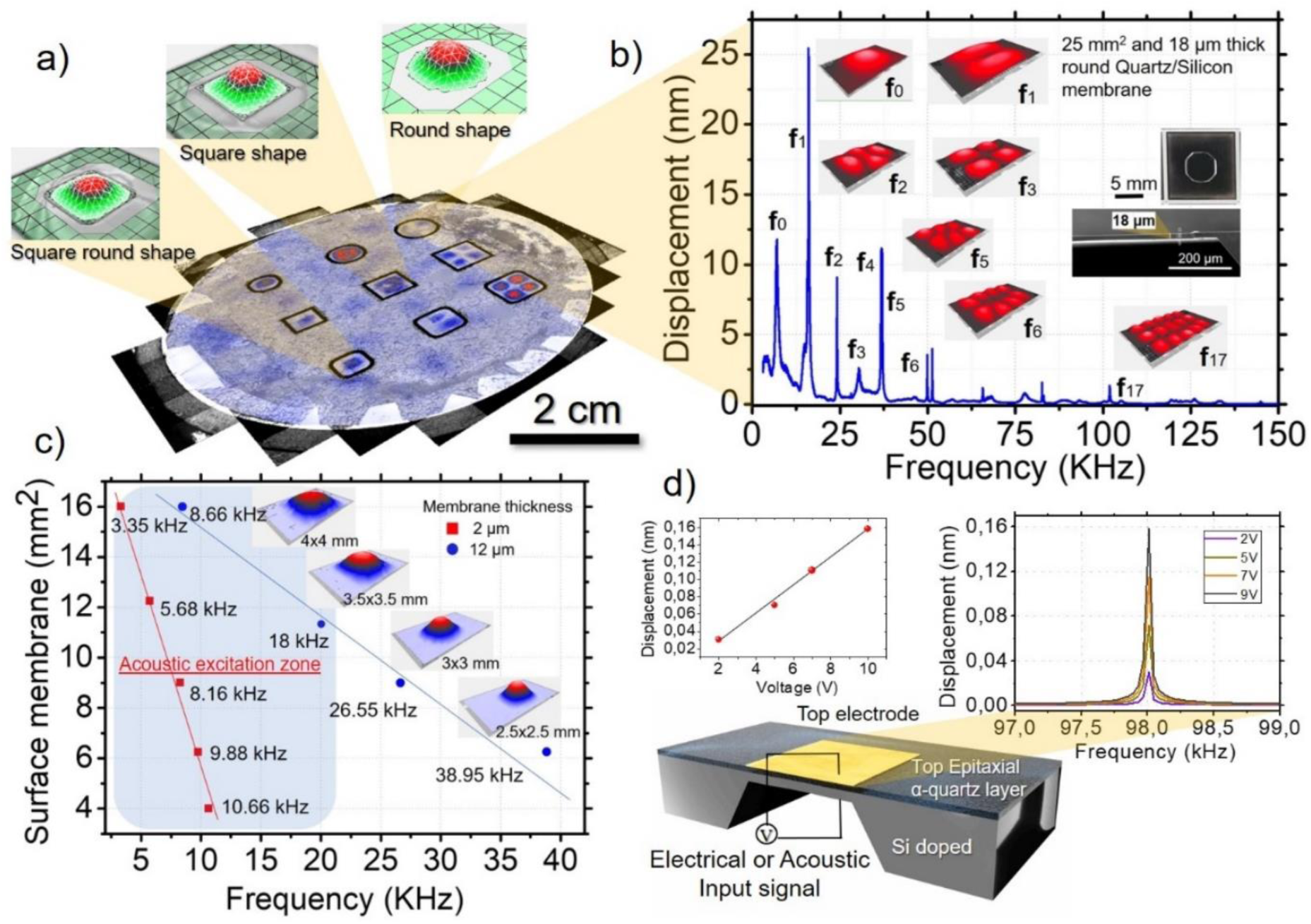
Microfabrication and electromechanical characterization of epitaxial α-quartz/Silicon-based MEMS resonators. **a)** Non-contact vibrometer measurements of a set of α-quartz/Silicon-based resonant membranes on 2 inches silicon wafer with control over resonant frequency, dimensions, thickness, and morphology. Notice that in the same α-quartz/Silicon wafer there are three different morphologies i.e. square, round and square round of 3, 4 and 5 mm of diameter or lateral dimension. **b)** Mechanical response of 25 mm^2^ and 18 µm thick round α-quartz/Silicon resonant membranes measured with an optical vibrometer excited using an acoustic input signal. **c)** Tailoring of the final resonance frequency of α-quartz/Silicon resonators trough the control of size and thickness of the resonator. Blue zone points out α-quartz/Silicon membrane resonators that can be activated with an acoustic input signal. **d)** Non-contact vibrometer measurements of a 1mm² piezoelectric α-quartz(100)/Si(100) membrane with a resonant frequency of 98 kHz activated by direct piezoelectric excitation using electrical input signal from 2 to 10 V). Notice that the membrane displacement amplitude is linear with the imposed voltage due to the inverse piezoelectric effect.

On the other hand, the top face of the device, i.e., the quartz active layer, is protected by screw-tightened wafer holders (also known as wafer chucks). We could control the laser etching process with high precision (see **Figure S9**) and finely finish the membrane fabrication by chemical means. As a result, this microfabrication process allowed us to preciously control the final thicknesses of high-quality quartz/Silicon MEMS resonators with a high mechanical quality factor and perfect vibrational motion as shown in **Figure 4b**. Moreover, this control over the size and thickness of the resonators made it possible the tailoring of the final resonance working frequency (see **Figure 4c**).

**Figure 4d** shows the successful piezoelectric activation of an Au/(100)quartz/doped-(100) silicon membrane vibrational motion through the indirect piezoelectric effect of the α-quartz active layer. We observed the linear behaviour between the input voltage signal and the displacement of the resonator, confirming the piezoelectric properties of quartz/silicon MEMS. However, the low piezoelectric coefficient of α-quartz (d_31_ = 2.4 pm/V)[14] and the small degree of bending of the membranes do not permit the complete piezoelectric transduction through the mechanical movements of quartz/silicon resonators. As a result, it was impossible to harvest piezo-generated charges from directly bending the resonators. Although it is possible to activate the movement of the resonators with the piezoelectric effect of the α-quartz layer, its displacement is low-performance, less than 1 nm (see **Figure 4d**). For this reason, we used membrane resonators with resonance frequencies in the acoustic zone, i.e., between 5 and 20 KHz (see **Figure 4c**), thus activating them acoustically using a microphone. This feature not only enabled us to study their mechanical behaviour in air and liquid mediums but also highlighted their potential for biosensing applications through optical transduction, inspiring our further research in the fabrication of a super high-frequency α-quartz NEMS membrane resonator with piezoelectric transduction at the intrinsic α-quartz material frequency, which depends on the thickness of the quartz.

## 6. Detection of Chikungunya virus particles with α-quartz/Silicon bio-MEMS

Next, we set out to perform mass measurements of the binding of different loads of CHIKV particles to α-quartz bio-MEMS in a liquid medium by non-contact vibrometry (**Figure 5a**). To render the bio-MEMS selective for CHIKV, we used a bioRL consisting of a layer of anti-CHIKV E2 biotinylated antibodies immobilized through the covalent surface conjugation of the SpyCatcher/SpyAvidin complex, as previously reported on α-quartz thin films (**Figure 3**). We equipped the Laser Doppler Vibrometer (LDV) with an environmental control chamber (temperature and % CO_2_) (**Figure S10**), and we performed the liquid measurements using a silicon well coupled to a microfluidic system for the real-time exchange of the different solutions. Using this configuration, we operated the bioMEMS sensor with a small sample volume of 25 μl. The setup allowed us to monitor the displacement in the frequency spectra of a squared α-quartz/silicon membrane of 25 mm^2^ and 18 μm thick during each step of the bio-conjugation process performed to obtain the bioRL (**Figure 5 b-c**). This included i) the covalent conjugation of a SpyCatcher layer at the surface of the piezo-membrane, ii) the subsequent covalent bonding of a SpyAvidin layer through the isopeptide bond formation between SpyCatcher and the SpyTag of the dead-streptavidin subunit and finally, iii) the immobilization of the anti-E2 biotinylated antibody with the single traptavidin subunit of the streptavidin tetramer of the SpyAvidin tetramer. As shown in **Figure 5b**, we observed a frequency signature of each layer of the bio-conjugation system, reaching ∼12 kHz upon the biotinylated antibody binding. The assembled α-quartz bioMEMS was then used to monitor the binding of CHIKV particles at different concentrations, from 6.13·10^9^ down to 3.06·10^6^ particles (**Figure 5c**). We found that increasing the number of viral particles per sample led to a systematic shift of the α-quartz membrane towards lower frequencies, indicating a mass response of the bioMEMS upon selective recognition of CHIKV loads.

**Figure 5.**
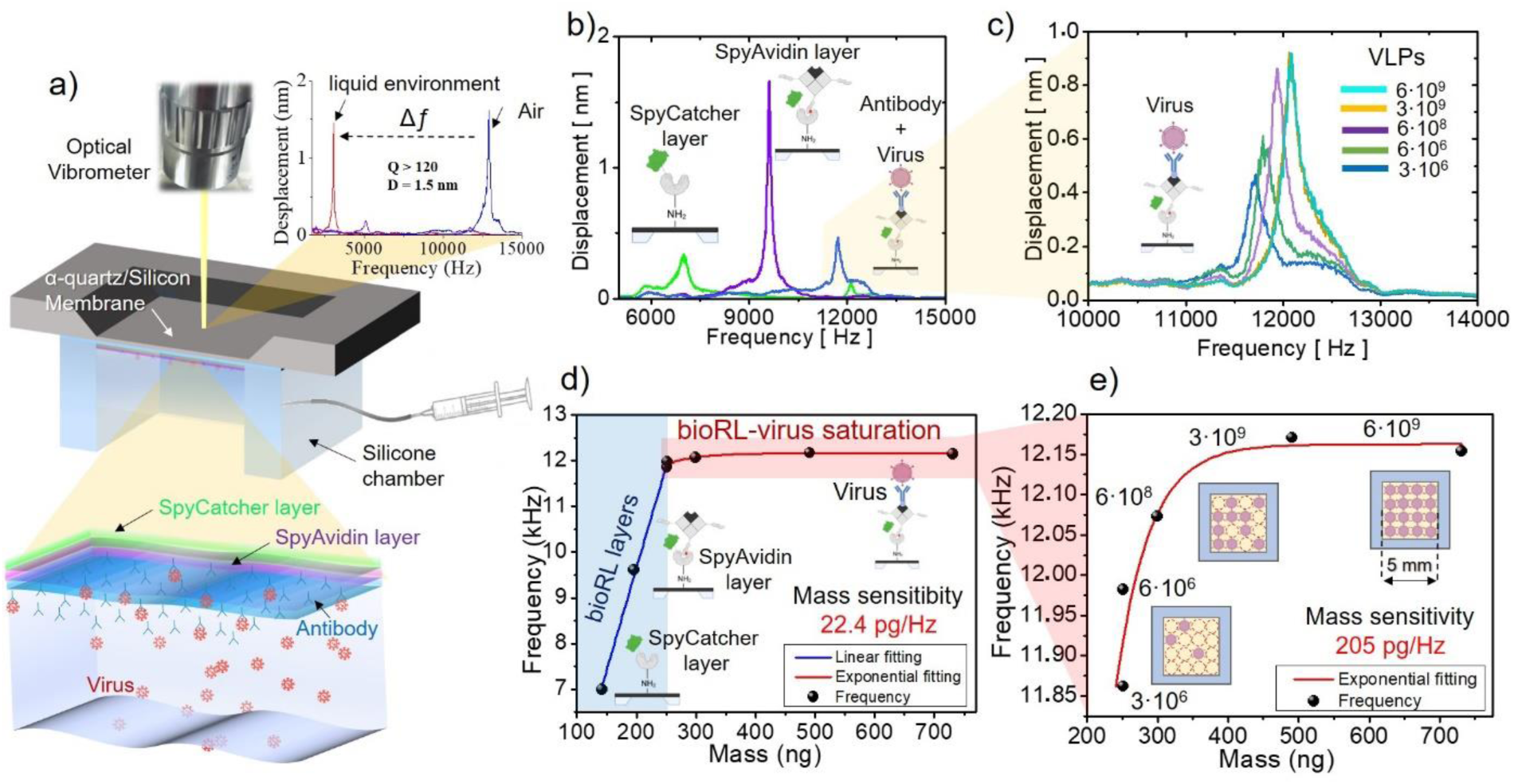
Non-contact vibrometry mass measurements of CHIKV binding to an α-quartz bio-MEMS functionalized with a virus bioRL. **a)** Schematic representation of the bioRL at the surface of the α-quartz membrane (consisting of a SpyCatcher layer, in green, followed by a SpyAvidin layer, in purple, and the biotinylated antibody, in blue, detecting CHIKV particles, in red) and the assembly of the α-quartz bio-MEMS device in liquid using a silicon-chamber microfluidic system to perform non-contact vibrometry measurements. **b)** Displacement (nm) versus frequency (Hz) spectra showing the signature of the different layers that constitute the bioRL at the surface of the α-quartz membrane: SpyCatcher layer (green), SpyAvidin layer (purple) and anti-E2 biotinylated antibody (blue). **c)** Displacement (nm) versus frequency (Hz) spectra of different CHIKV loads (from 3·10^6^ to 6·10^9^ virus like particles, VLPs) captured by bioRL of the α-quartz membrane. **d)** Relationship between the frequency (Hz) and mass change (ng) of each layer of the bioRL and the charged CHIKV loads captured by the bioRL fitted by a linear and exponential saturation model respectively. **e)** Magnification of the relationship between the frequency (Hz) and mass change (ng) of the captured CHIKV loads captured fitted by an exponential saturation model.

To assess the mechanical sensitivity of our α-quartz/silicon membrane resonator, we measured the resonance frequencies associated with the masses of each protein layer in the bio-RL (as detailed in the experimental section), as well as the virus captured on the membrane.

To this end, we fitted the frequency versus mass data to a linear model for the different bioRL and used an exponential saturation model for the CHIKV particles (see Figure 5d). From the slope of the linear model, we determined the sensitivity of the α-quartz/Silicon membrane resonator to be 22.4 pg/Hz (see, blue region of the plot in **Figure 5d**). Furthermore, we obtained an acceptable accordance between the theoretical and measured mass of the different layers of the bioRL quantified with the α-quartz bio-MEMS, as indicated by a mean residual value of ∼ 2.13 ng.

To establish the virus detection limit of the bioRL due to the saturation of its available binding sites assuming a maximum occupancy of the membrane surface, we tested different viral loads ranging from 3.06·10^6^ to 6.12·10^9^ CHIKV particles and measured the corresponding resonance frequencies (see **Figure 5e**). The concentration of CHIKV particles in the samples was established by light scattering using a nanoparticle tracking analyser (**see Figure S11**). Then, the theoretical mass was calculated using the buoyant density of 1.23 g/mL of CHIKV particles, assuming that the viral particles are roughly spherical with a diameter of ∼60 nm[32]. The data were then fitted using an exponential saturation model (see equation 2 in the experimental section) allowing to precise estimation of the CHIKV particle’s occupancy on the α-quartz/Silicon membrane surface (See **Figure 5e**). Consequently, we could establish the membrane sensitivity from the derivative at the point of intersection with the linear model, yielding a value of 205 pg/Hz (**see** **Figure 5e**). Notably, the similar order of magnitude obtained for the sensitivity values derived from the linear and saturation models suggest a consistency between the two approaches used.

Furthermore, from these experiments we could establish an experimental limit of detection (LoD) of the CHIKV-selective bioMEMS of 9 ng/mL, i.e., 3 x 10^8^ viral particles/ml, corresponding to a mass of approximately 0.23 ng (as detailed in the experimental section). This LoD is nearly 5 times more sensitive than the lowest LoD reported using an ELISA assay targeting viral samples (50 ng/ml)[32] but, still significantly higher than the RT-PCR detection limit of 0.08 pg[33]. Therefore, this indicates that our LoD is well within the detectable range expected for benchmarked diagnosis methods. Indeed, the obtained detection level should be suitable for various diagnostic and research applications, as it falls within the range of sensitivity for both immunological and molecular detection methods commonly used for virus detection. Furthermore, it provides a convenient balance between sensitivity and practicality for most laboratory procedures.

## 7. Towards a high-frequency piezo-NEMS based on α-quartz membranes

After demonstrating the potential of bio-conjugated epitaxial α-quartz/silicon membranes as highly sensitive biosensors for CHIKV virus detection, we explored the possibility of fabricating α-quartz NEMS resonators with piezoelectric transduction at the intrinsic α-quartz material frequency. Therefore, paving the way towards reaching compact biosensors with even higher sensitivities and implementing these performances on a portable commercial device.

To this aim, we fabricated a super high frequency α-quartz NEMS capable to resonates at the intrinsic piezoelectric α-quartz thin film frequency by completely removing the silicon substrate (see **Figure 6a**). These super high resonance frequency resonators consisted of 2D α-quartz nanometric membranes of 120 nm thick with a gold electrode set in a sandwich (see **Figure 6a**). To fabricate this type of high frequency resonators it was necessary to completely remove the silicon to avoid unwanted physical interface phenomena that prevent a perfect electrical isolation of the quartz layer. We developed a modified version of our previous etching process (see **Figure S12**) to stabilize the 2D α-quartz membrane over large surfaces. In that case, we used a poly(methyl methacrylate) PMMA transparent thin film to both electrically isolate the top electrode and mechanically stabilize the thin quartz membrane (see **Figures 6b-e** and Figure S8). To remove the silicon substrate, we changed the etching agent from TMAH to KOH allowing us to make the process more environmentally friendly, safer and faster, and avoiding the need to use a clean room.

**Figure 6.**
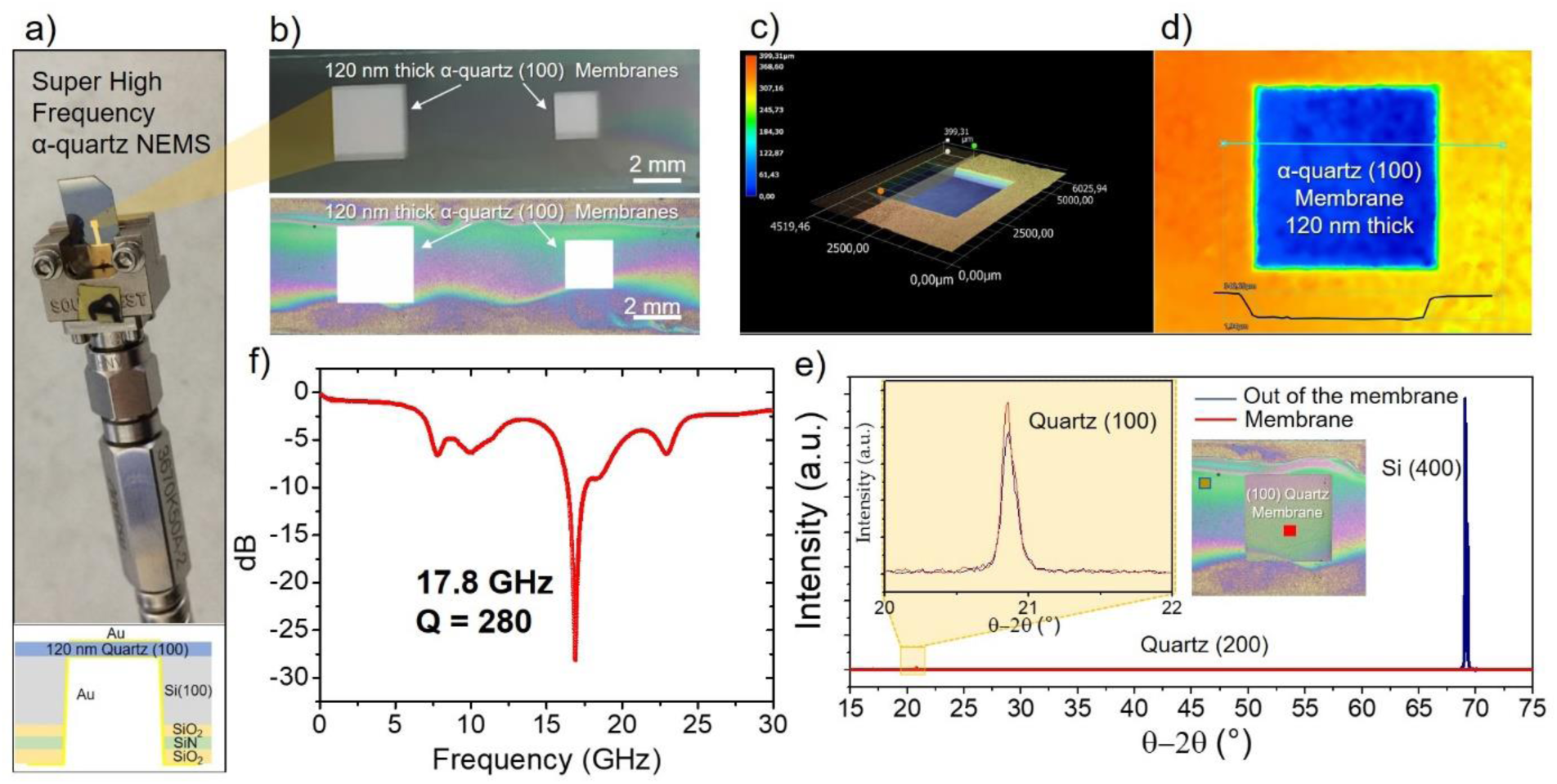
Piezoelectric resonance at super high frequency of 120 nm thick quartz membrane resonators. **a)** Image of the final (100) α-quartz NEMS consisted of a squared shape 120 nm thick membrane with 3 mm of lateral dimension. Below, schematics showing the gold electrode set in sandwich form where the whole of the silicon substrate has been removed, **b)** Optical images with different transmission lighting showing the perfect transparency of the 120 nm thick quartz membrane. Notice that the white color of the transmitted light confirms the absence of silicon since the silicon absorbs red light (see Figure S1). **c)** 3D optical image of the quartz nanomembrane and its in-depth profile. **d)** Two-dimensional optical heat mapping showing the depth of the silicon substrate etching up to the quartz layer and the perfect planarity of the resonator. **e)** XRD diffraction measurement in and out of the epitaxial quartz membrane confirming the total removal of the silicon substrate. **f)** Resonance frequency at 17.8 GHz with reflection values of -28.04 dB, respectively analyzed using a vector network analyzer. This result confirms the resonant behavior of the α-quartz layer, indicative of its piezoelectric properties.

We measured the resonance frequency of quartz NEMS by using a vector network analyzer and we observed that the epitaxial (100) quartz thin film resonates at 17.8 GHz (see **Figure 6f**). This is to our knowledge, the first nanometric 2D α-quartz crystal with super high resonance frequency. The experimental resonant value of 17.8 GHz fully corresponds to our theoretical calculations which depends on the transverse velocity of sound propagation in α-quartz i.e. 3460 m/s (see equation 3 in the experimental section). Hence, if we consider an α-quartz layer of about 100 nm the resonant frequency should be around 17.3 GHz. More importantly, we calculated an experimental quality factor of 280 at air at 300K in (100) quartz NEMS, which is in the same order of quartz bulk. Notice that the maximum Q for a high stability quartz oscillator can be estimated as Q = 1 × 10^13^ / f, where f is the resonance frequency in Hertz. As a result, the maximum Q for a quartz bulk at 17.3 GHz is around 500 which is the same order than the epitaxial (100) quartz films NEMS. This α-quartz NEMS technology combined with the developed bioconjugation system for selective virus detection (**Figure 3** and **5**) might overcome the current technological bottlenecks and, importantly, the limited working frequencies and sensitivity of the current quartz transducers made from bulk micromachining or hybrid integration methods, making it possible to engineer the first piezoelectric transductor PoC device enhanced mass resolution for arbovirus detection.

## 8. Conclusions

In conclusion, we have combined a cost-effective and wafer-scale integration of piezoelectric (100) α-quartz thin films on Si (100) using chemical methods, addressing a long-standing challenge to synthesizing high-quality quartz thin films on an industrial scale. Using chemical processes and thermal treatment, we have been able to control the nucleation, crystallization, microstructure, and thicknesses of epitaxial α-quartz films on silicon up to 4 inches while preserving a coherent (100)quartz/(100)silicon crystalline interface. As a result, we could achieve the fabrication of wafer-scale piezoelectric MEMS and high-quality factor NEMS resonators at super high-frequency, from 5 KHz to 17.8 GHz. These novel quartz-based devices offer significant advantages over bulk quartz in terms of size, power consumption, integration cost, and ability to provide multiple operation frequencies. In this direction, we showed that by developing a versatile bioconjugation strategy based on the ultrastable SpyCatcher/SpyAvidin complex combined with commercial biotinylated antibodies, it is possible to render piezoelectric bioMEMS selective for the recognition of arboviruses, such as the CHIKV, over other viral loads. Using non-contact vibrometry coupled to a microfluidics system we showed that quartz bioMEMS can reach a mass sensitivity of 22.4 pg/Hz in liquid conditions with a virus detection limit of 9 ng/ml, which is 5 times more sensitive than conventional ELISA tests targeting virus samples[32]. Collectively, this work opens the door towards low-cost and miniaturized single-chip solutions for high-frequency and ultra-sensitive mass devices that could be used as PoC testing devices in the biomedical sector but also find applications in any field of the electronics and sensing devices sector.

## 9. Experimental Section

### Sol gel preparation and wafer scale deposition and growth of epitaxial (100) quartz on silicon

All chemicals were obtained from Sigma-Aldrich and used as received without further purification. Solution A was prepared by dissolving 1 g of Brij-58 (CAS number 64-17-5) in 23.26 g of absolute ethanol (CAS number 9004-95-9), followed by the addition of 1.5 g of HCl (37%) (CAS number 7647-01-0) and 4.22 g of tetraethyl orthosilicate (TEOS) (CAS number 78-10-4). The solution was stirred for 18 hours. A 2M aqueous solution of Sr²⁺ (solution B) was prepared using SrCl₂·6H₂O (CAS number 10025-70-4). Solution C, used for the deposition of Sr-silica films via spin coating, was prepared by adding 275 μL of solution B to 10 mL of solution A, followed by stirring for 10 minutes. Films were deposited within 40 minutes of preparing solution C to avoid instability of Sr²⁺. The Sr/SiO₂ molar ratio in solution C was 0.05, with a final molar composition of TEOS:Brij-58:HCl:EtOH:SrCl₂ = 1:0.44:0.7:25:0.1.

Gel films on Si(100) substrates (until 4 inch) were prepared using a vacuum-free Ossila spin coater. During the dip-coating process, the ambient temperature and relative humidity were set to 25 °C and 40%, respectively, and the deposition conditions were set at a spinning speed of 2000 rpm for 30 seconds. After deposition, films were consolidated with a thermal treatment at 450 °C for 5 minutes in air atmosphere. The multilayer gel films were obtained by repeating several times the process of mono-layer preparation on the same substrate.

As-prepared gel films were introduced into a furnace already at 1000 °C in an air atmosphere and held at this temperature for 300 min. The crystallized films were recovered after cooling down the furnace to room temperature.

### Electron, X-ray diffraction and optical analysis of epitaxial quartz wafers

Bright field images in STEM mode were acquired with a Nion UltraSTEM operated at 100 kV. A JEOL F200 transmission electron microscope (TEM) ColdFEG operated at 200kV was used for the structural characterization of the interface via electron diffraction. A lamella prepared by Focused Ion Beam (FIB) in a ThermoFisher Scios 2 was inserted in a JEOL analytical tomography holder for three-dimensional acquisitions. The TEM was used in STEM mode and STEM images (512 x 512 pixels) were recorded from a JEOL high-angle annular dark-field (HAADF) detector in Gatan Digital Micrograph. Electron diffraction patterns were acquired with a Gatan OneView camera, a CMOS-based and optical fibre-coupled detector of 4096 by 4096 pixels. The electron beam in STEM mode was aligned following a self-made protocol[34] so that the most quasi-parallel condition could be fulfilled with a 120-nm beam diameter (Probe size 8 and 10-µm condenser aperture). The acquisition of the 3D ED data that enabled the reconstruction of the diffraction spaces was carried out with the (S)TEM-ADT module [35], a self-developed Digital Micrograph plug-in freely available at https://github.com/sergiPlana/TEMEDtools. This software automatizes the acquisition of 3D ED datasets by automatically inserting and retracting the HAADF detector, setting the diffraction and imaging conditions of the electron beam, and collecting the STEM reference images and diffraction patterns. The program follows the methodology initially implemented by, but it is modified to allows the acquisition of more than one diffraction pattern per tilt angle at any desired positions of the reference image[36]. In this work, two patterns were acquired per tilt angle (at quartz and silicon) for an angular range between -40° and 40° of alpha-tilt angle of the sample holder with a tilt step of 1° (162 diffraction patterns in total). Nominal camera length was set to 500 mm, and a precession angle of 1.15° was used for each diffraction pattern through a P2000 prototype precession unit by Nanomegas SPRL. PETS2 software[37] was used for unit cell determination and data reduction, and Jana2020[38] for dynamical refinements through the dyngo module[39].

Atomic STM resolution images were performed by using a FEI Titan3 operated at 80 kV and equipped with a superTwin® objective lens and a CETCOR Cs-objective corrector from CEOS Company.

The crystalline textures, rocking curve measurements, and epitaxial relationship of quartz films and cantilevers were performed on a Bruker D8 Discover diffractometer equipped with a 2D X-ray detector (3 s acquisition each 0.02° in Bragg–Brentano geometry, with a radiation wavelength of 0.154056 nm). 2D and 3D optical images of quartz films and membranes resonators were obtained in a Keyence® VHX7000 optical digital microscope. The microstructures of the quartz films and MEMS were investigated with a FEG–SEM model Su-70 Hitachi, equipped with an EDX detector X-max 50 mm^2^ from Oxford Instruments. The topography of quartz films was studied by AFM in a Park Systems NX-Scanning Probe Microscopy unit.

### Vibrometry Measurements at air and liquid cell conditions

The 3D vibration reconstructed images and spectra of the fabricated quartz/silicon MEMS were evaluated with a laser doppler vibrometer LDV equipped with laser, photodetector, and frequency generator. The vibrometer (MSA-600, Polytech®) allowed characterizing in-plane and out-of-plane motion through non-contact silicon encapsulation, without the need to prepare or decapsulate the device up to 25 MHz. Then, the dynamic displacement of the resonating MEMS sensor is reconstituted by the PSV software.This equipment has an internal frequency generator utilized to actuate the inverse piezoelectricity of quartz membranes.

### Quartz/silicon membrane MEMS manufacturing

A 2-inch (100)Si wafer coated with the thin piezoelectric epitaxial quartz film. The wafer was opened using a laser to create its moving part. The power and repetition rate of which varies according to the thickness of the wafer and the desired depth of the moving part with different morphologies and size. The 2-inch silicon wafer is then placed in a support from AMMT to protect the thin film of quartz before being immersed in a TMAH bath heated to 80°C by water bath. A protective layer is applied by sputtering (PECVD) to the wafer surface, but on the other side from the part covered with quartz. This protective layer is composed of three successive deposition: silicon nitride (400nm) silicon dioxyde (400 nm) and again silicon nitride (400 nm). Then, the protective layer is etched until the silicon bulk is revealed ether by ICP-RIE with a CHF_3_/O_2_ plasma or with the laser etching system. If the ICP-RIE is chosen for the etching of the protective layer, a lithography step, before the RIE etching, is mandatory in order to have the desire a desired membrane’s design. Then, a laser beam of 3.75 μm in diameter and 5 W in power is used to etch the membrane by uploading a designed membrane file in laser machine. Due to this flexible and easy etching method, we can etch any shape of membrane we want in a few minutes. Before the TMAH (CAS number 75-59-2) etching step the native oxide layer that is formed on the silicon surface is removed by using Hydrofluoric acid (BOE 10%, CAS number 76664-39-3).

### Estimation of the recognition layers and virus mass and modeling

To estimate the mass of each layer of the bioRL, i.e. SpyCatcher, SpyAvidin complex, and anti-E2 biotinylated antibody layer, we assumed spherical volumes with diameters ranging from 4 to 15 nm, considering their corresponding molar masses and crystal structure data [40,41]. SpyCatcher-GFP protein has an average molecular weight of ∼ 43kDa, considering the weight of SpyCatcher to be 16kDa [40] and 27kDa for the GFP tag, and an approximative dimension of ∼ 4 nm measured from PDB 4MLI crystal structure. SpyAvidin hub protein used (Tre1DTag3) has molecular weight of ∼ 65 kDa[29,41] and an approximative dimension of ∼ 5-8 nm, based on the PDB 2Y3E crystal structure of the apo Traptavidin homo tetramer.

For the monoclonal biotinylated IgG2b antibody layer, we estimated a molecular weight of an IgG antibody of ∼ 150 kDa and an approximative dimension of ∼ 12-15 nm, based on the PDB 1IGT crystal structure[42]. Biotinylation, which involves attaching biotin molecules to the antibody adds a mass of 0.24 kDa, which might be negligible compared to the antibody’s overall mass. Therefore, we estimated that the mass of a monoclonal biotinylated antibody is approximately 150 kDa, equivalent to about 2.49 × 10^19^ grams per molecule.

To ensure complete coating of the 5 x 5 mm squared membrane resonator in preparation for the recognition layer, we used an excess of proteins. The mass of the proteins covering the membrane resonator was calculated using the following formula

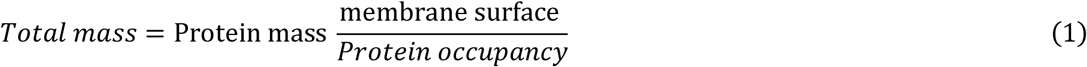

The frequency versus mass data were fitted to a linear model. From the slope of this model we obtained the sensitivity of the membrane, i.e., 22.4 pg/Hz with a mean residual value of ∼ 2.13 ng.

An exponential saturation model was used to analyze the viral loads ranging from 3.06·10^6^ to 6.12·10^9^ virus particles, which were experimentally measured using light scattering (see figure S11). These measurements together with the corresponding resonance frequencies allowed to determine the virus detection limit based on the maximum virus occupancy expected on the 25 mm^2^ membrane resonator surface. Subsequently, we calculated the theoretical mass using the buoyant density of 1.23 g/mL of CHIKV particles, assuming that the particles are roughly spherical with a diameter of 60 nm[32]. Then, the data was fitted according to the exponential saturation model:

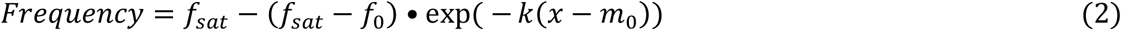

Where *f_sat_* represents the saturation frequency, *f*_0_ is the resonance frequency without any attached viruses, *m*_0_ is the mass of the recognition layer without any attached viruses, and *k* is the saturation coefficient.

The sensitivity of the membrane in this model was estimated from the derivative at the point of intersection with the previous linear model, obtaining a value of 205 pg/Hz.

### (100) Quartz membrane NEMS manufacturing

To obtain a quartz membrane, first, samples are cleaned in acetone/ethanol, then several successive layers of PE-CVD consisting of Si_3_N_4_/SiO_2_/Si_3_N_4_ (500 nm per layer) are deposited on the back side to create an etching mask. The lithography is performed on the protectives layers in order to achieve the wanted design for the membrane. An Az1518 photoresist is spin coated at 3000 rpm for 30 seconds then baked at 110°C for 1 min. A 20s exposure is done by using an UV lamp through a “stencil” type mask and the sample is dipped into AZ726 developer for 1 min. The protection is then etched by plasma ICP-RIE CHF_3_/O_2_ to create the openings until the silicon is exposed. The silicon is then chemically etched (KOH 45%) from the back to the quartz layer. The front face is protected by a made-to-measure shield. As the quartz layer is composed of SiO_2_, it acts as a barrier layer for the KOH etching. Because the quartz layer is thin (i.e., 150 nm) they become fragile during KOH etching and the resulting membranes tend to collapse on themselves. To avoid this problem, a 300 nm-thick layer of PMMA resin is deposited directly on the quartz surface (front face of the sample) using a spinner (3000 rpm, 30 sec) and then a hard bake at 140°C for 1 min before KOH etching. The resin maintains the quartz structure and is resistant to the etching solution for a given amount of time. The etching speed of silicon in KOH 45% at 80°C is 1 µm·min^-1^. The result is a transparent film of quartz and PMMA. The resin can be further removed using an O_2_ plasma.

### High frequency piezoelectric experimental measurement and theoretical calculations of 120 nm thick epitaxial quartz membranes NEMS

An Anritsu 37397D vector network analyzer was used to characterize a layer of α-quartz deposited on silicon. The analyzer, covering a frequency range from 40 MHz to 65 GHz, was configured to measure the S11 parameter, representing the reflection magnitude at the input port. The experimental setup was designed as follow: The α-quartz of silicon sample was connected to port 1 via a Southwest connector, ensuring low-loss measurement integrity. A SOLT (Short, Open, Load, Thru) calibration was performed to eliminate extraneous reflections and establish a reference point for accurate S11 measurements. The S11 parameter was measured in the forward reflection mode over a frequency range of 0.04 to 40 GHz. The results, shown in **Figure 6**, display resonance minima at approximately 17.8 GHz with reflection values of -28.04 dB, respectively. These dips confirm the resonant behavior of the α-quartz layer, indicative of its piezoelectric properties.

The theoretical calculation of the resonance frequency for a α-quartz crystal depends on the transverse velocity of sound propagation in α-quartz which has a value of 3460 m/s. If we consider a quartz layer of about 100 nm the resonant frequency should be around 17.3 GHz which is of the same order as the experimentally obtained frequency (see equation 6.).

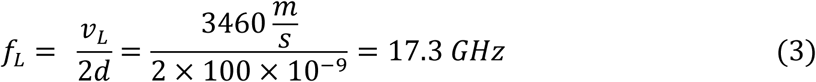

Importantly, the experimental quality factor of up to 280 at air obtained in (100) quartz NEMS is the same order of quartz bulk since the maximum Q for a high stability quartz oscillator can be estimated as Q = 1.6 × 10^7^ / f, where f is the resonance frequency in megahertz. As a result, the maximum Q at 17.3 GHz is around 500 which is the same order than the epitaxial (100) films quartz NEMS.

### SpyCatcher and SpyTag purification

SpyTag003-mKate2 (Addgene ID 133452) and SpyCatcher003-sfGFP (Addgene ID 133449) were a gift from Mark Howarth, previously reported in[26].

Briefly, to purify SpyTag-mKate2 and SpyCatcher-GFP, *E. coli* BL21 CodonPlus (DE3) RIPL (Agilent Technologies) cells were transformed with plasmid pET28a-SpyTag003-mKate2 or pJ404-SpyCatcher003-sfGFP. Cells expressing SpyTag003-mKate2 were grown in the autoinduction medium (Formedia) supplemented with 50 µg/mL of kanamycin and 34 µg/mL of chloramphenicol at 30 °C for 24h with shaking at 200 rpm. Cells expressing SpyCatcher003-GFP were grown in LB containing 50 µg/mL of ampicillin, 34 µg/mL of chloramphenicol and 0.8% glucose. After reaching OD_600_ = 0.5-0.6, the cultures were inoculated with 0.42 mM IPTG and incubated at 30 °C with shaking at 200 rpm for 4-5 h.

For SpyTag-mKate2 and SpyCatcher-GFP purifications, cells were harvested and lysed by sonication in lysis buffer (50 mM Tris-HCl pH 7.8, 300 mM NaCl) supplemented with a cocktail of protease inhibitors (complete EDTA-free, Roche), 1 mM PMSF, and 1mg/ml lysozyme. Clarified cell lysates were centrifuged at 30,000 *g* for 30 min at 4°C before purification using Ni-NTA resin (Qiagen) according to the manufacturer’s instructions. After elution, proteins were dialyzed into 20 mM Tris-HCl pH 8. Proteins were finally concentrated through Amicon Ultra filter units (10 kDa cut-off), quantified using a Bradford assay (Biorad) and flash-frozen in liquid N_2_ and stored at 80°C.

### Purification of SpyAvidin Hubs

Traptavidin-E6 (Addgene ID 59549) and Dead Streptavidin-SpyTag (Addgene ID 59548), previously described in[29], were a gift from Mark Howarth.

Purification of SpyAvidin Hubs from Dead-Streptavidin-SpyTag (DTag) and Traptavidin-E6 (Tre) was performed as described in[29]. Briefly, RIPL cells (Agilent) were transformed with 50 ng of pET21-DTag and pET21-Tre. The transformed cells were plated on LB agar containing ampicillin and incubated overnight at 37°C. Cells were grown from a single colony for 5 hours at 37°C in 1 L of LB medium with ampicillin and protein expression was induced by adding 100 µg/mL IPTG, followed by 4 hours of incubation at 37°C. Cells were harvested by centrifugation at 4500 rpm for 15 minutes at 4°C, and the pellets were stored at -80°C for subsequent purification. The next purification steps were handled at 4°C. For inclusion body purification, pellets were resuspended in 10 mL PBS containing EDTA-free protease inhibitors, 1 mM PMSF, 1% Triton X-100, 10 mM EDTA, and 0.1 mg/mL lysozyme, and incubated on ice for 1 hour. The suspensions were sonicated on ice with four cycles of 40-second pulses at 40% intensity, with 2-minute rest intervals between cycles. Cell debris were removed by centrifugation at 10,000g for 30 minutes at 4°C. The pellets were washed with PBS containing 1% Triton X-100 and 10 mM EDTA, followed by another wash with PBS and 10 mM EDTA. The inclusion bodies were resuspended in 6 M guanidine hydrochloride (pH 1.0) on ice for 1 hour and centrifuged again at 10,000g for 30 minutes at 4°C to separate the supernatant containing solubilized inclusion bodies. The *A*_280_ of the dissolved inclusion bodies was then measured and an equivalent number of absorbance units of the appropriate variant was mixed together: 5,9 mL of Tre dissolved inclusion bodies with an *A*_280_ of 37,8 were mixed with 4,2 mL of DTag dissolved inclusion bodies with an *A*_280_ of 53,3. Refolding of the proteins in tetramers was achieved by slow dilution into 200 mL of PBS containing 10 mM EDTA at 4°C, stirred overnight. Ammonium sulfate (60 g) was added to the refolded protein solution, stirred for 1 hour at 4°C, and filtered. Another 60 g of ammonium sulfate was added, and the solution was centrifuged at 15,000g for 30 minutes at 4°C to collect the protein pellet. The pellet was resuspended in 50 mL of sodium borate buffer (50 mM sodium borate, 300 mM NaCl (pH 11.0), and centrifuged at 10,000g for 30 minutes at 4°C. The clarified supernatant was purified using a 5 mL iminobiotin-sepharose affinity column, equilibrated with sodium borate buffer. The proteins were eluted with 20 mM KH_2_PO_4_ (pH 2.2), and eluate was dialyzed against 20 mM Tris-HCl (pH 8.0) at 4°C. The eluate was then exchanged into 20 mM Tris HCl pH 8.0 by dialysis and loaded onto a 5 mL Q-HP column (GE Healthcare). The different forms were isolated by using 50 column volumes (250 mL) linear gradient of 0.15–0.4 M NaCl and collecting 5 mL fractions with a 2 mL/min flow rate using 20 mM Tris·HCl pH 8.0 as the running buffer. Fractions of 1.5 mL were collected. Fractions containing tetramers with 1 Traptavidin-E6 and 3 Dead-Streptavidin-SpyTag proteins were pooled and analyzed by SDS-PAGE to confirm purity and integrity. The purified tetramer was concentrated using a Vivaspin concentrator with a 30 kDa cutoff, and protein concentration was measured at A_280_. The final protein solution was dialyzed against PBS, aliquoted, flash-frozen in liquid N_2_ and stored at -80°C.

### Bio-conjugation of α-quartz MEMS and thin films

3-aminopropyldimethylethoxysilane (APDMES) was from abcr. Acetic acid, hydrochloric acid, and ethanol (EtOH) were purchased from Sigma-Aldrich. Maleimide-Malhex-NHS was from Biopharma PEG.

α-quartz thin films and membranes were thoroughly washed with acetic acid and for 45 minutes in a glass petri dish. Then, samples were rinsed five times with ultrapure water. Cleaned samples were placed into a plasma cleaned for 5 min. Immediately after the samples were immersed in a freshly prepared solution of APDMES:EtOH (1:20) mixture for 10 min at room temperature. Next, samples were rinsed with EtOH and then with ultrapure water. All samples were gently blow dried under N_2_ and subsequently put in an oven at 80°C for 30 minutes by covering their surface against dust. After this step, silicon chambers (ibidi) were placed on the surface of α-quartz thin films and membranes and a solution of sodium borate was added for 1 hour at room temperature. Samples were then dried under N_2_ and subsequently incubated with Maleimide-Malhex-NHS (MMN, at 5mM diluted in sodium borate) for 1 hour at room temperature. Samples were rinsed five times with phosphate buffered saline (PBS pH = 7.4) solution. SpyCatcher-GFP diluted in PBS at the desired concentration was incubated for 1 hour at room temperature. After thoroughly rinsing the surface, they were incubated overnight with a bovine serum albumin (BSA) solution at 1% in PBS to block the SpyCatcher-GFP conjugated layer. Next, samples were rinsed in PBS and SpyTag-mKate2 or SpyAvidin hubs at the desired concentration were added for 1h at room temperature. Finally, samples were rinsed and analyzed under a microscope (airyscan/confocal or MSA 600 doopler vibrometer system).

### Production of pseudo-typed viral particles

To produce CHIKV-like particles, subconfluent HEK 293T cells were seeded in 10 cm dishes and transfected with 30 µg of C-Env-Ruby3 plasmid by the calcium phosphate precipitation method. The medium was changed 6 h post-transfection and the supernatant was collected 72 h post-transfection. The supernatant was clarified by low speed centrifugation for 10 minutes, filtered through a 0.2 µm membrane filter and pelleted through a 20% sucrose cushion in Hepes 20mM pH 7.4, NaCl 150mM, EDTA 0.1mM buffer at 120,000g for 2 h at 4°C using a Beckman SW32-Ti rotor. The pellet was gently resuspended in Hepes 20mM pH 7.4, NaCl 150mM, EDTA 0.1mM buffer and CHIKV-like particles were kept at 4°C until use.

To produce VSV-G-like particles, subconfluent HEK 293T cells were seeded in 10 cm dishes and transfected by the calcium phosphate precipitation method with 30 µg of the following plasmid DNAs: pRRL-SFFV, psPax2, pCDNA3.3-Gag-mCherry, and pMD.G (VSV-G) at a ratio of 2:1:1:0,5 respectively as described in [43]. The medium was changed 6 h post-transfection and the supernatant was collected 72 h post-transfection. This supernatant was clarified by low speed centrifugation for 10 minutes, filtered through a 0.45 µm and pelleted through a 20% sucrose cushion in Hepes 20mM pH 7.4, NaCl 150mM, EDTA 0.1mM buffer at 120,000g for 2 hours at 4°C using a Beckman SW32-Ti rotor. The pellet was gently resuspended in Hepes 20mM pH 7.4, NaCl 150mM, EDTA 0.1mM buffer and VSV-G-like particles were kept at 4°C until use.

After each preparation, the concentration of pseudo-typed viral particles was established by light scattering with a nanoparticle tracking analyzer (ZetaView® x30 series, Particle Metrix).

C-Env-Ruby3 plasmid was generated as described by André-Arpin *et al*. (unpublished results from L. Picas’ lab, IRIM, Montpellier). pRRL-SFFV was a kind gift from C. Goujon’s lab, IRIM, Montpellier, and psPax2 and pCDNA3.3-Gag-were a kind gift from R. Gaudin’s lab, IRIM, Montpellier,

### Airyscan/confocal microscopy and image analysis

Images were acquired on a Zeiss LSM980 Airyscan/confocal Microscope (MRI facility, Montpellier). Excitation sources were 405 nm diode laser, an argon laser for 488 and 514 nm, and a helium/neon laser for 633 nm. Acquisitions were performed on a 63X/1.4 objective. Multidimensional acquisitions were acquired via an Airyscan detector (32-channel GaAsP photomultiplier tube array detector).

Airyscan/confocal images were quantified using the mean grey value of the different fluorescence intensities by ImageJ[44].

### Data representation and statistical analysis

Data representation was performed using Origin and Prism GraphPad software. Statistical analysis was performed using the Mann-Whitney two-tailed test or ordinary one-way ANOVA multiple comparisons test, with single pooled variance using Prism GraphPad software. In all statistical, the levels of significance were defined as: *P<0.05, **P<0.01, ***P<0.001 and ****P<0.0001.

## Supporting Information

Supporting Information is available from the Wiley Online Library or from the author.

## Acknowledgements

This project had received funding from the European Research Council (ERC) under the European Union’s Horizon 2020 research and innovation programme (project SENSiSOFT No.803004 and Project QOVID No. 101082079). This project has received financial support from the CNRS through the MITI interdisciplinary programs through its exploratory research program. We also acknowledge financial support from Projects No. PID2020-118479RBI00 and PID2023-152225NB-I00. The authors are grateful to Félix Rico and Claire Valoteau for helpful discussions. The authors thanks Fatima El Alaoui, Jérome Feuillard and Mickael Blaise at IRIM for assistance in protein purification. The authors thank D. Montero for performing the FEG–SEM images and chemical analysis. The FEG–SEM instrumentation was facilitated by the Institut des Matériaux de Paris Centre (Grant No. IMPC FR2482). The authors thank Phillipe Nouvel and HERMES technical platform, for the expertise and advice during the high frequency measurements. The authors thank Frederic Pichot, for the expertise and advice during the membranes lithographic processes at the CTM platform. This work has been supported by the Labex NUMEV to R.R., A.C-G and L.P. We acknowledge the imaging facility MRI, member of the national infrastructure France-BioImaging supported by the French National Research Agency (ANR-10-INBS-04, «Investments for the future»). Electron microscopy observations at ORNL were supported by the Office of Science, Materials Sciences and Engineering Division of the U.S. Department of Energy.

## Notes

### Competing Interest Statement

The authors have declared no competing interest.

